# Striatal Dopamine Can Enhance Learning, Both Fast and Slow, and Also Make it Cheaper

**DOI:** 10.1101/2024.02.14.580392

**Authors:** Andrew Westbrook, Ruben van den Bosch, Lieke Hofmans, Danae Papadopetraki, Jessica I. Määttä, Anne GE Collins, Michael J. Frank, Roshan Cools

**Author notes:** **Materials & Correspondence** Correspondence should be addressed to Andrew Westbrook at.

## Abstract

Associations can be learned incrementally, via reinforcement learning (RL), or stored instantly in working memory (WM). While WM is fast, it is also capacity-limited and effortful. Striatal dopamine may promote RL plasticity, and WM, by facilitating updating and effort exertion. Yet, prior studies have failed to distinguish between dopamine’s effects on RL versus WM. N = 100 participants completed a paradigm isolating these systems in a double-blind study measuring dopamine synthesis with [^18^F]-FDOPA imaging and manipulating dopamine with methylphenidate and sulpiride. Learning is enhanced among high synthesis capacity individuals and by methylphenidate, but impaired by sulpiride. Methylphenidate also blunts effort cost learning. Computational modeling reveals that individuals with high dopamine synthesis rely more on WM, while methylphenidate boosts their RL rates. The D2 antagonist sulpiride reduces accuracy due to diminished WM involvement and faster WM decay. We conclude that dopamine enhances both slow RL, and fast WM, by promoting plasticity and reducing effort sensitivity. These results also highlight the need to control for dopamine’s effects on WM when testing its effects on RL.

## Introduction

Striatal dopamine signaling has been implicated in cortico-striatal plasticity [1–3] and reinforcement learning (RL) across species [2,4]. Yet, in humans, a substantial contribution to learning is mediated by working memory (WM) processes [5–12] and prior work linking individual differences in dopamine function to RL typically fails to disentangle the impact on incremental RL versus WM.

While WM is faster and more flexible, multiple constraints limit its contributions. Unlike RL, WM is capacity-limited and subject to decay [13–15]. WM may thus play a smaller role when there are more items to remember, and when they were encountered farther in the past. WM is also effort-costly, and people may forgo WM-based strategies to avoid the effort [16–19].

Importantly, theoretical models and empirical data implicate striatal dopamine signaling in both RL, via alterations of synaptic plasticity, and WM via formation and expression of cortico-striatal synapses that encode gating policies [20–23]. Furthermore, striatal dopamine can also make WM less effort-costly [24–28]. While WM is effortful, striatal dopamine signaling promotes willingness to exert effort, by making people more sensitive to potential benefits, and less sensitive to effort costs [24]. If striatal dopamine signaling promotes reliance on WM - either by shaping corticostriatal synapses to facilitate gating, or by making WM less effortful - this can amplify effective learning rates, confounding inferences about the direct effects of striatal dopamine signaling on RL.

Here, we use a paradigm designed to distinguish between contributions of WM and RL to stimulus-response learning. The Reinforcement Learning Working Memory (RLWM) task [10] manipulates the degree to which people can rely on WM. Set sizes vary (between two and five items) across blocks of trials, thus taxing WM load, delay, and interference to varying degrees. While healthy adults *can* rely mostly on WM (rather than RL) when there are only two items to encode recent experience, participants *must* rely increasingly on RL as set sizes grow to exceed WM capacity [7,9,10,29]. We furthermore include a surprise test phase, after learning, to probe the durability of RL-informed learning in terms of participants’ ability to recall features of the stimuli after a long delay.

To study dopamine’s effects on RL and WM, we employ a combination of methods. We isolate the effects of dopamine signaling in the striatum by measuring individual differences in the rate at which dopamine is synthesized in presynaptic striatal terminals using [^18^F]-FDOPA PET imaging. We also manipulate dopamine signaling, while participants perform the RLWM task in separate sessions, by administering placebo, methylphenidate – a dopamine (and noradrenaline) reuptake blocker commonly used to treat attention deficit hyperactivity disorder, or sulpiride – a D2 receptor antagonist commonly used to treat psychosis in schizophrenia.

To anticipate our results, we find that striatal dopamine is related to enhanced performance on the RLWM task. Behavioral and computational modeling reveals that higher dopamine synthesis capacity promotes faster learning by increasing reliance on WM. Sulpiride undermines performance specifically because it increases interference within WM when there is an increasing number of items in between successive encounters of the same stimulus. Furthermore, we find that methylphenidate boosts the rate of RL, controlling for WM contributions to the learning process. Finally, we find that while WM is effort-costly, methylphenidate can also blunt implicit effort cost learning associated with more demanding tasks.

## Results

In each of three drug sessions – placebo, methylphenidate, and sulpiride – participants completed the Reinforcement Learning Working Memory (RLWM) task, which was designed to dissociate RL and WM processes during stimulus-response learning [10]. On a given trial, participants learn to associate pictures with one of three buttons by trial-and-error (Figure 1). Participants are given feedback about the accuracy of each response. To distinguish contributions of RL versus WM, stimuli are presented in blocks of varying set size. By this design, WM can dominate when there are only two items to remember, typically separated by minimal delays (e.g., one or two trials). Conversely, RL necessarily plays a larger role for larger set sized blocks (up to five items in a block) as delays and the number of items grow to exceed WM capacity.

**Figure 1.**
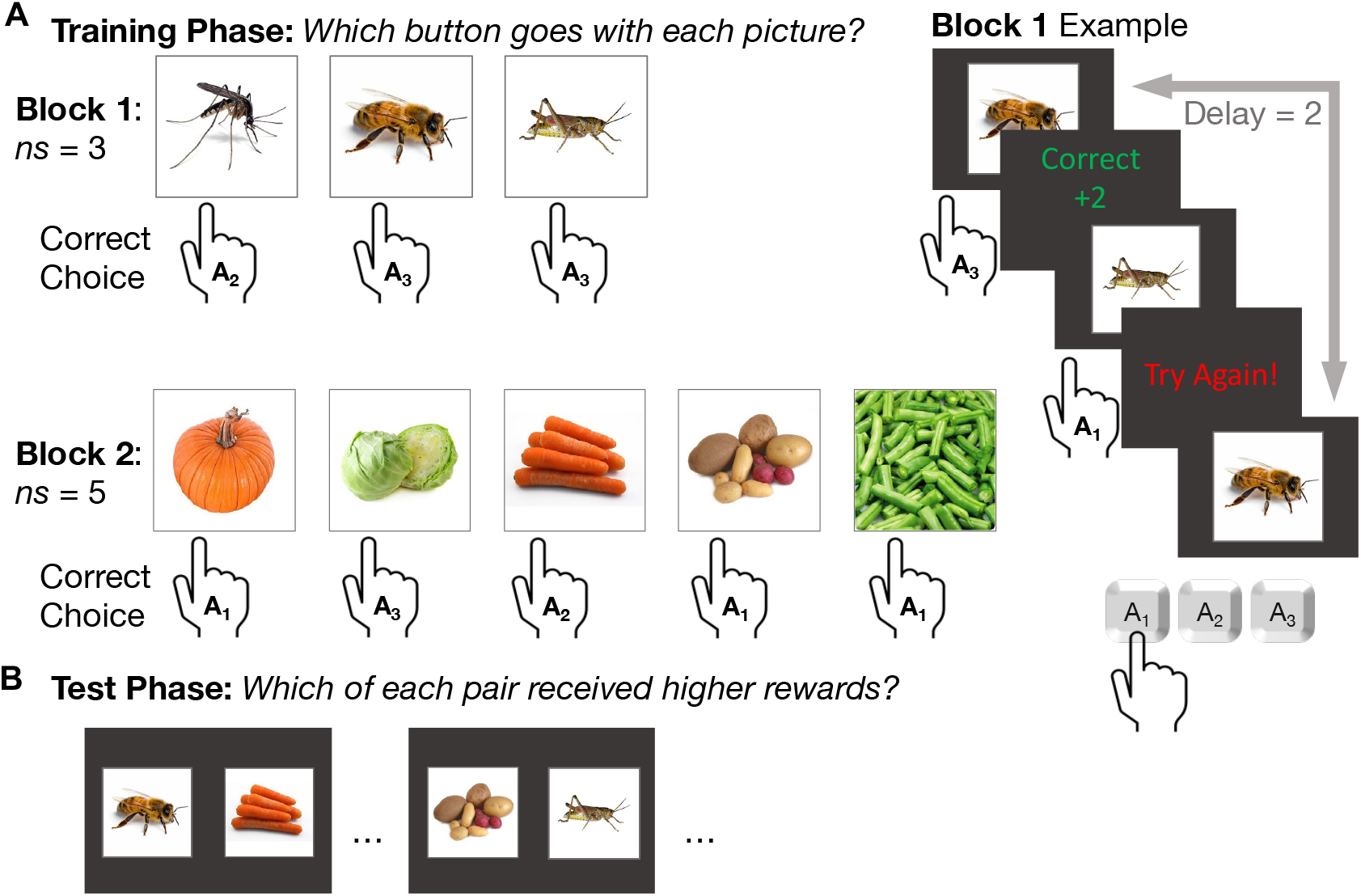
RLWM task schematic. In each block participants are shown sets of images with set size (*ns*) varying by block (between *ns* = 2 and *ns* = 5). Participants learn through trial and error which of three buttons to press for each stimulus (A_j_). If they respond correctly, they are rewarded points (+2 or +1, probabilistically, see Methods for full details) and if they are incorrect, they receive no points.

### Both reinforcement learning and working memory contribute to performance

Conjoint contributions of WM and RL are implied by the shape of learning curves by stimulus iteration (Figure 2). Curves are steeper when set size is smaller and people could, in principle, rely more on WM. In a logistic regression of accuracy on set size and previous iteration count, accuracy is higher with more iterations of each stimulus (β = 1.90; p < 2.0×10^−16^) and for smaller set sizes (β = -.30; p < 3.6×10^−10^). An interaction between these factors implies that effective learning rates are larger in smaller set size blocks (β = -.11; p = .0016). Prior computational and neurophysiological work [7,9,10,29,30] support the hypothesis that steeper learning curves for smaller set sizes reflects greater reliance on WM rather than faster RL.

**Figure 2.**
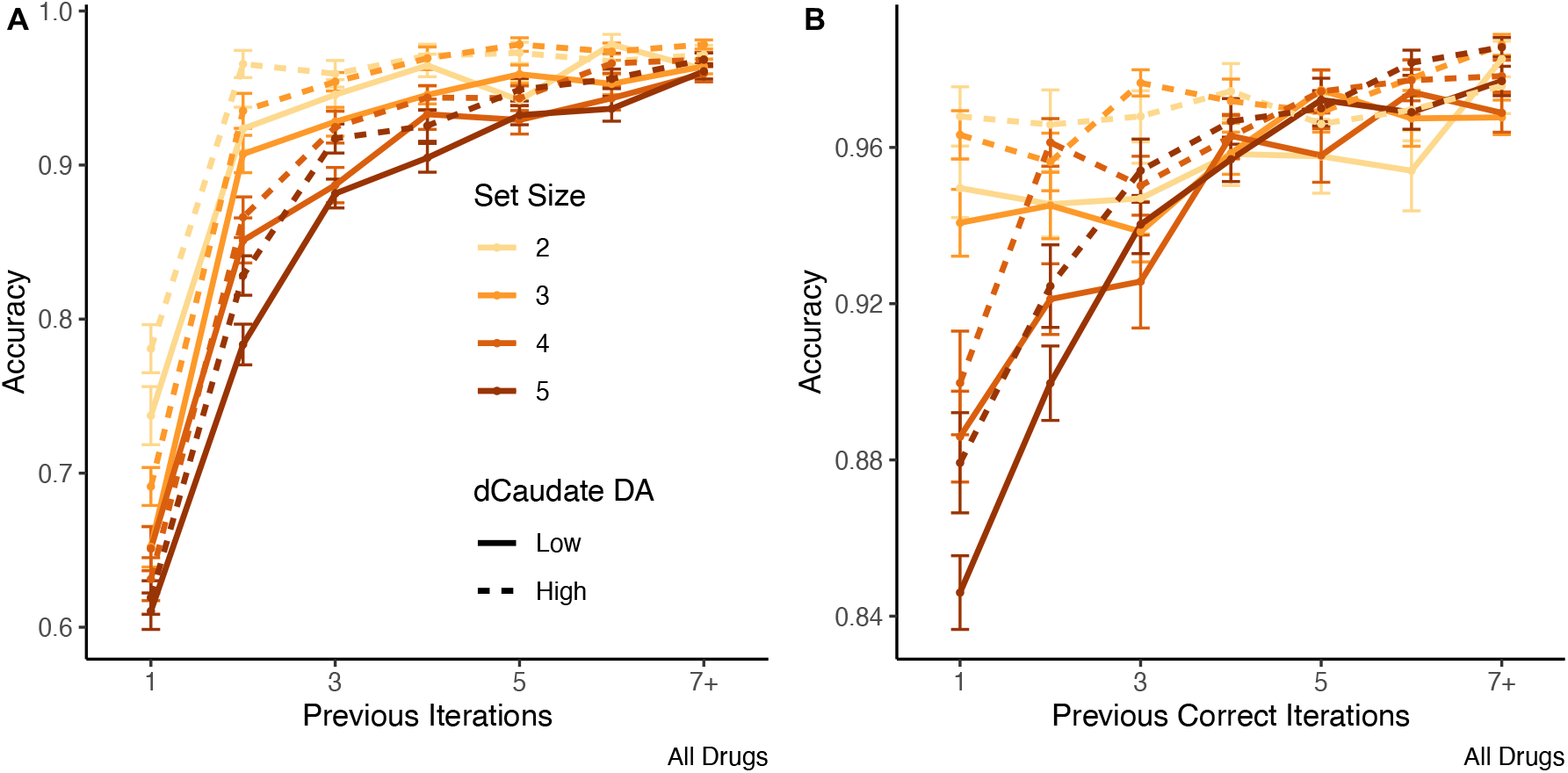
Accuracy as a function of set size, individual differences (median split) in dopamine synthesis capacity in the dorsal caudate nucleus (dCaudate DA), and **A**) the number of previous iterations for each stimulus or **B**) the number of previous correct iterations for each stimulus. Accuracy increases with more iterations of each stimulus on all drugs, and it is higher for those with higher dopamine synthesis capacity.

It has further been demonstrated in both behavior and neural activity that people rely more on WM for novel items when fast and flexible updating offers the best performance benefits, and they rely more on RL for familiar items when slow but robust RL-cached values have had the chance to form [9,10,29,30]. An analysis of performance early versus late in a block supports this distinction, with WM appearing to contribute more to performance early, while RL contributes more late in a block. Two hallmarks of WM reliance - sensitivity to set size (Figure 3A) and delay (Figure 3B) - are apparent early in a block (fewer than three previous correct iterations for each stimulus) and are minimal late in a block (the last two iterations for each stimulus) across all three drug sessions (Figure 3). Two-way ANOVAs reveal that participants’ average performance across all three sessions varies as a function of set size (F_1,732_ = 127, p < 2.0×10^−16^), early versus late in a block (F_1,732_ = 7.28×10^3^, p < 2.0×10^−16^) and their interaction (F_1,732_ = 104, p < 2.0×10^−16^), and also effects of delay (F_1,2388_ = 596, p < 2.0×10^−16^), early versus late (F_1,2388_ = 102, p < 2.0×10^−16^), and their interaction (F_1,2388_ = 30.5, p = 3.6×10^−8^). Thus, these data support the hypothesis that when people encounter novel items, they rely heavily on WM, and thus also show limitations due to capacity and interference within WM, but they shift to RL as the number of previous correct iterations grows and RL-based information becomes more reliable. This inference is also consistent with evidence that although people perform better for smaller set sizes early in a block, neural indices of RL grow across a block and do so faster in higher set sized blocks, when they rely more on RL versus WM [30].

**Figure 3.**
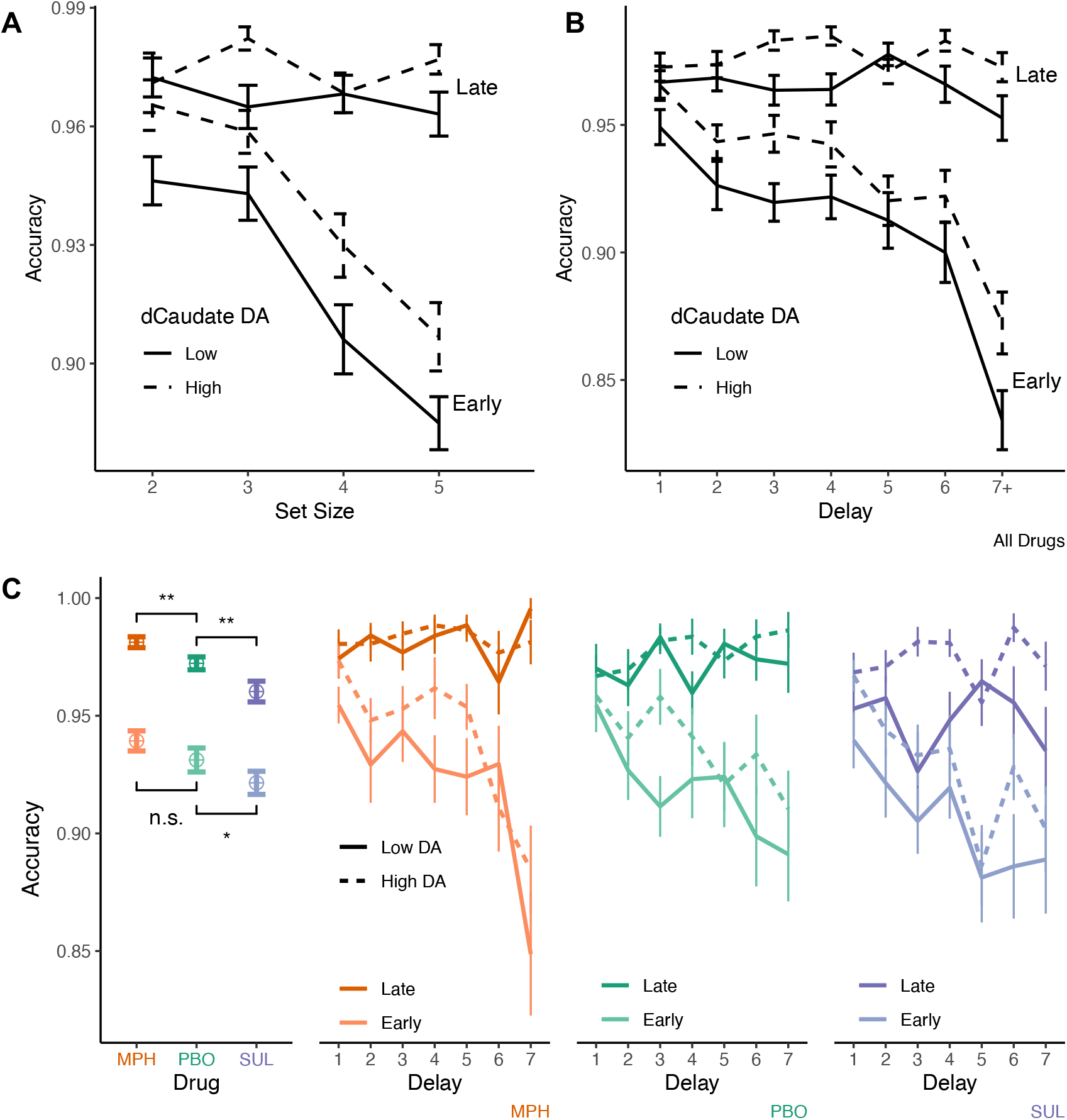
Accuracy as a function of set size, delay, and dopamine synthesis capacity in the dorsal caudate nucleus (dCaudate DA) across all three drug sessions: methylphenidate (MPH), placebo (PBO), and sulpiride (SUL), early (the first two iterations of all stimuli) versus late (the last two iterations) in each block. The effects of **A)** set size and **B)** delay are evident early, but not late in each block. **C)** Means and standard errors reflect that methylphenidate significantly increases and sulpiride decreases accuracy relative to placebo both early and late in a block (albeit the increase in early performance on methylphenidate is non-significant). Further analyses reveal that methylphenidate boosts performance in late versus early trials to a greater extent than placebo (*,** indicate p < .05 and p < .005 in mixed effects logistic regressions of accuracy; described in the next section “*Striatal dopamine variously enhances performance*”).

### Striatal dopamine variously enhances performance

Striatal dopamine signaling improves performance in multiple ways. We employed complementary tools to dissect these effects. First, we measured individual differences in dopamine synthesis capacity using [^18^F]-FDOPA PET imaging. Dopamine synthesis capacity is correlated across five striatal sub-regions (ICC = 0.75; p = 3.1×10^−4^), yet we focused our analysis on the dorsal caudate nucleus, where dopamine signaling has previously been implicated in reinforcement learning about higher cognition [31–34], working memory gating [20,23,26,35], and cost-benefit decision-making about cognitive effort [24,36–38]. We also had participants complete the task in three drug sessions, after taking methylphenidate – a dopamine and noradrenaline transporter blocker which should amplify striatal dopamine signaling, and sulpiride – a selective D2 receptor antagonist, or placebo.

To evaluate the effect of dopamine on performance, we fit a hierarchical Bayesian regression of trial-wise accuracy on individual differences in dopamine synthesis capacity and drug, controlling for session number. In our model we also simultaneously estimate the effects of set size, the number of previous correct iterations, and the delay since the last correct iteration, along with higher order interactions with drug status and dopamine synthesis capacity (see Supplementary Table S1 for the full results). The fitted model reveals that striatal dopamine signaling clearly enhances performance. Specifically, higher dopamine synthesis capacity (β = .17; p = .026) and methylphenidate versus placebo (β = .41; p = 1.6×10^−6^) both increase accuracy, while sulpiride decreases accuracy (β = -.27; p = 1.7×10^−4^; Figure 3).

To understand why striatal dopamine synthesis capacity and methylphenidate boosted performance and sulpiride undermined it, we fit a reinforcement learning model to behavior (adapted from [10,12]) examining how people learn to select the correct action for each stimulus in each block. The algorithm combines a WM component featuring instantaneous learning, capacity limits, and susceptibility to decay, and a capacity-unlimited RL component with incremental learning rates (see Methods for full details).

Fitted model parameters imply two ways in which striatal dopamine signaling boosts performance. First, it boosts performance by increasing the likelihood that people rely on WM (*ρ*; Figure 4A,B), which facilitates fast and flexible acquisition of new associations. Second, striatal dopamine signaling also appears to increase the learning rate in the RL system (*α*_*RL*_; Figure 4C) which describes the rate at which incrementally acquired stimulus-response associations can contribute to action selection.

**Figure 4.**
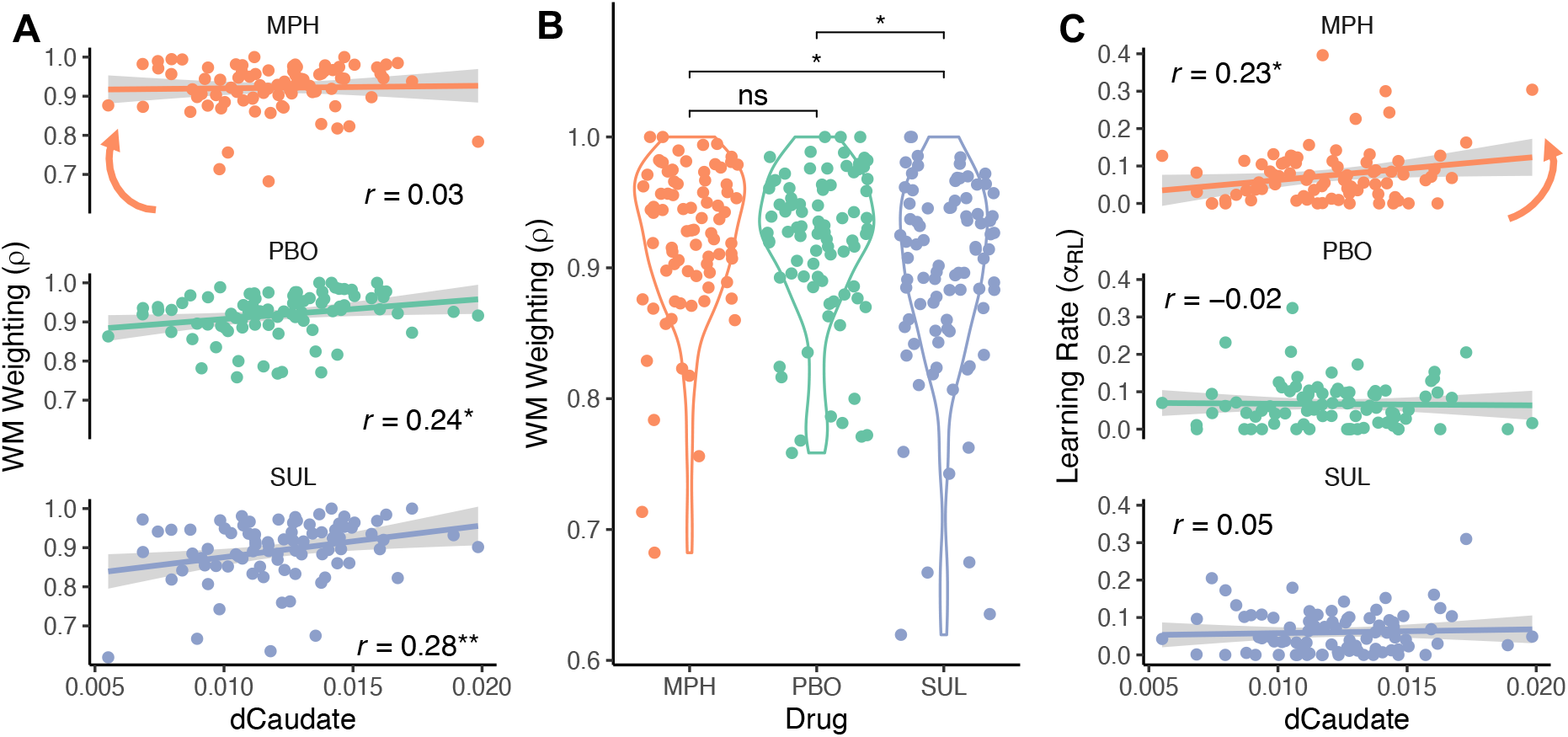
**A**,**B)** WM weighting (*ρ*) and **C)** the RL learning rate (*α*_*RL*_) parameters from reinforcement learning algorithm as a function of individual differences in the dopamine synthesis capacity in the dorsal caudate nucleus of the striatum across three drug sessions. Drug effects and Spearman’s correlation values are reported along with their significance level (*,** indicate p < .05 and p < .005).

A hierarchical regression of the parameter *ρ* (WM reliance) on dorsal caudate dopamine synthesis capacity and drug, controlling for session number, reveals an overall positive effect of dopamine synthesis capacity across sessions (β = .23; p = .029; Figure 4A). This correlation supports the hypothesis that people who synthesize dopamine at a higher rate rely more on WM in general. There was also a negative effect of sulpiride versus placebo (β = -.35; p = .0086; Figure 4B) indicating that WM contributed less to choice, on sulpiride. Methylphenidate did not affect the baseline propensity to use working memory (β = .083; p = .48). However, there was a trending negative interaction between baseline dopamine synthesis capacity (β = -.20; p = .078) suggesting that methylphenidate tends to increase reliance on WM more for those who synthesize dopamine at a lower rate (Figure 4A).

The hypothesis that participants with higher dopamine synthesis capacity rely more on WM is further supported by evidence that they tend to perform better early in a block when items are novel and WM affords better performance (Figure 3A,B). A hierarchical logistic regression, restricted to early trials, regressing accuracy on set size, drug, and dopamine synthesis capacity reveals that participants with higher dopamine synthesis capacity perform better when WM plays a bigger role (β = .14; p = .023). In contrast, the same model fitted to late trials reveals a diminished effect of dopamine synthesis capacity (β = .12; p = .17). Moreover, there is also evidence linking dopamine synthesis capacity to WM reliance in the full logistic regression (including all trials and task variables; Supplementary Table S1). Namely, participants with higher dopamine synthesis capacity have a larger effect of set size on accuracy (β = -.10; p = .029). Additionally, a two-way interaction indicating stronger delay effects in larger set sized blocks (β = -.28; p = 2.8×10^−14^) is also larger for participants with a higher dopamine synthesis capacity (β = -.08; p = .016). Thus, model-independent analyses converge on the hypothesis that people who synthesize dopamine at a higher rate rely more on WM, thus boosting their performance, especially on early trials, and also making them more susceptible to set size and delay effects on trial-wise accuracy.

Even when controlling for the effect of striatal dopamine signaling on WM, however, there is evidence that dopamine boosts incremental RL learning rates, as noted above (Figure 4C). A hierarchical regression of the learning rate parameter *α*_*RL*_ on dopamine, controlling for session number, reveals a two-way interaction such that methylphenidate boosts learning rates for those with higher dopamine synthesis capacity (β = .30; p = .032). There were no main effects of dopamine synthesis capacity (β = -.027; p = .79), methylphenidate (β = .15; p = .29), or sulpiride (β = -.10; p = .46). However, the interaction resulted in a positive correlation between dopamine synthesis capacity and *α*_*RL*_ during the methylphenidate session (*r* = .23; p = .042). These results support the hypothesis that by blocking dopamine reuptake and amplifying post-synaptic signaling, methylphenidate increases the rate of reinforcement learning, especially for those who synthesis dopamine at a higher rate.

The inference that methylphenidate boosts learning rates is also supported by model-independent analyses. Specifically, the full logistic regression of accuracy on dopamine and task variables (Supplementary Table S1) reveals that methylphenidate increases accuracy (β = .41; p = 1.6×10^−6^) and amplifies the effect of previous correct iterations (two-way interaction: β = .20; p = 3.7×10^−3^; Figure 5) – a basic index of RL processes [9,30,39]. Although dopamine synthesis capacity also increases accuracy overall (noted above), it does not, in contrast with methylphenidate, interact with previous correct iterations (β = -.01; p = .84). Thus, on methylphenidate, people perform better in part because their incremental improvement with each rewarded trial is greater. Consequently, the overall improvement in accuracy between early and late trials is larger on methylphenidate versus placebo. A mixed-effects logistic regression of accuracy restricted to just early and late trials indeed reveals that the improvement between early and late trials is larger on methylphenidate versus placebo (two-way interaction: β = .48; p = .0044). These results converge on the hypothesis that methylphenidate accelerates RL, leading to faster incremental acquisition of stimulus-response contingencies, and a bigger overall improvement from early to late trials in each block.

**Figure 5.**
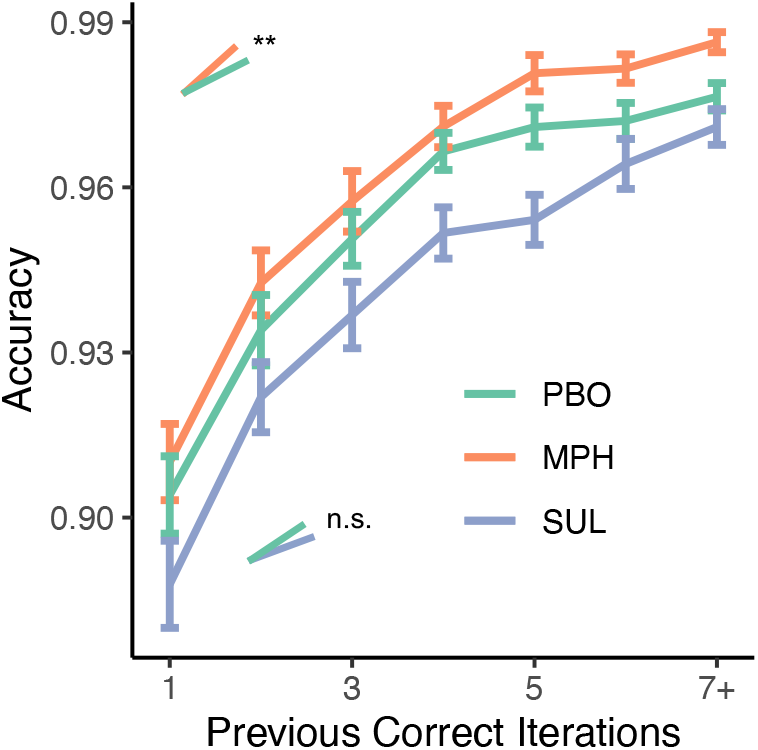
Accuracy as a function of previous correct iterations for each stimulus and drug. The effect of previous correct iteration is larger on methylphenidate (MPH) versus placebo (PBO), and no different on sulpiride (SUL; ** indicates p < .005).

Sulpiride, in contrast, undermines performance. According to mixed-effects logistic regressions of accuracy, restricted to either early or late trials, this is true both early (β = -.21; p = .027) and late in a block (β = -.42; p = .0038). One reason that people performed worse on sulpiride appears to be that they rely less on working memory, in general. Indeed, fitted parameters from the reinforcement learning algorithm reveal that sulpiride had a negative main effect on the WM reliance term *ρ* (β = -.35; p = .0028), as noted above. There was no interaction between sulpiride and dopamine synthesis capacity, however (β = .11; p = .36). Thus, just like on placebo, the positive correlation between *ρ* and dopamine synthesis capacity on sulpiride remained (β = .28; p = .0010).

There is evidence that people relied less on WM overall, after taking sulpiride, because WM itself was less reliable. In the full logistic regression of accuracy on dopamine and task variables (Supplementary Table S1), the negative effect of sulpiride on accuracy, was accompanied by a two-way interaction indicating that delay effects were worse on sulpiride versus placebo (β = -.11; p = .047). Thus, participants tend to perform worse when there is a bigger delay since the last previous correct iteration for a given stimulus, to a greater extent on sulpiride versus placebo. Thus, sulpiride may have reduced performance, in part, because it amplifies delay effects, undermining the contributions of WM to the learning process.

In sum, parameters from our reinforcement learning model converge with our model-independent analyses on the interpretation that greater striatal dopamine signaling among those who synthesize dopamine at a higher rate predicts greater reliance on working memory. However, when dopamine signaling is boosted by methylphenidate, blocking dopamine reuptake from the synaptic cleft, those with low dopamine synthesis capacity tend to rely more on working memory, matching high dopamine synthesis capacity participants. Conversely, on sulpiride, performance decreases and this appears to reflect less reliance on working memory across participants. Our results furthermore indicate that methylphenidate increases the rate of reinforcement learning, especially among those who synthesize dopamine at a higher rate, even after controlling for the drug’s effects on working memory.

### Methylphenidate blunts cognitive effort discounting

Following stimulus-response learning, participants are presented with a surprise test phase in which they are asked to select which of two stimuli received greater rewards (Figure 1). Stimuli were selected pseudo-randomly from those previously encountered across all training phase blocks. The intent of this test phase was to study RL-based value representations – which should be robust to the decay across blocks – after learning. Following prior work [7,30,39], we examine whether the reward statistics people learn through RL are influenced by the degree to which people discount rewards by the cognitive effort they exerted when rewards were received.

To evaluate the effects of task and dopamine variables on RL-based cached values, we fit a hierarchical Bayesian logistic regression of pairwise accuracy (correctly identifying the stimulus which received higher rewards) on task variables, drug, and dopamine synthesis capacity (see Supplementary Table S2 for full results). Replicating prior work [39] we find that participants faithfully track reward statistics, correctly picking the stimulus which was rewarded at a higher rate (intercept: β = .08; p = .0090), and increasingly so as the difference in actual rewards increased (β = .31; p = 4.2×10^−15^; Figure 6A).

**Figure 6.**
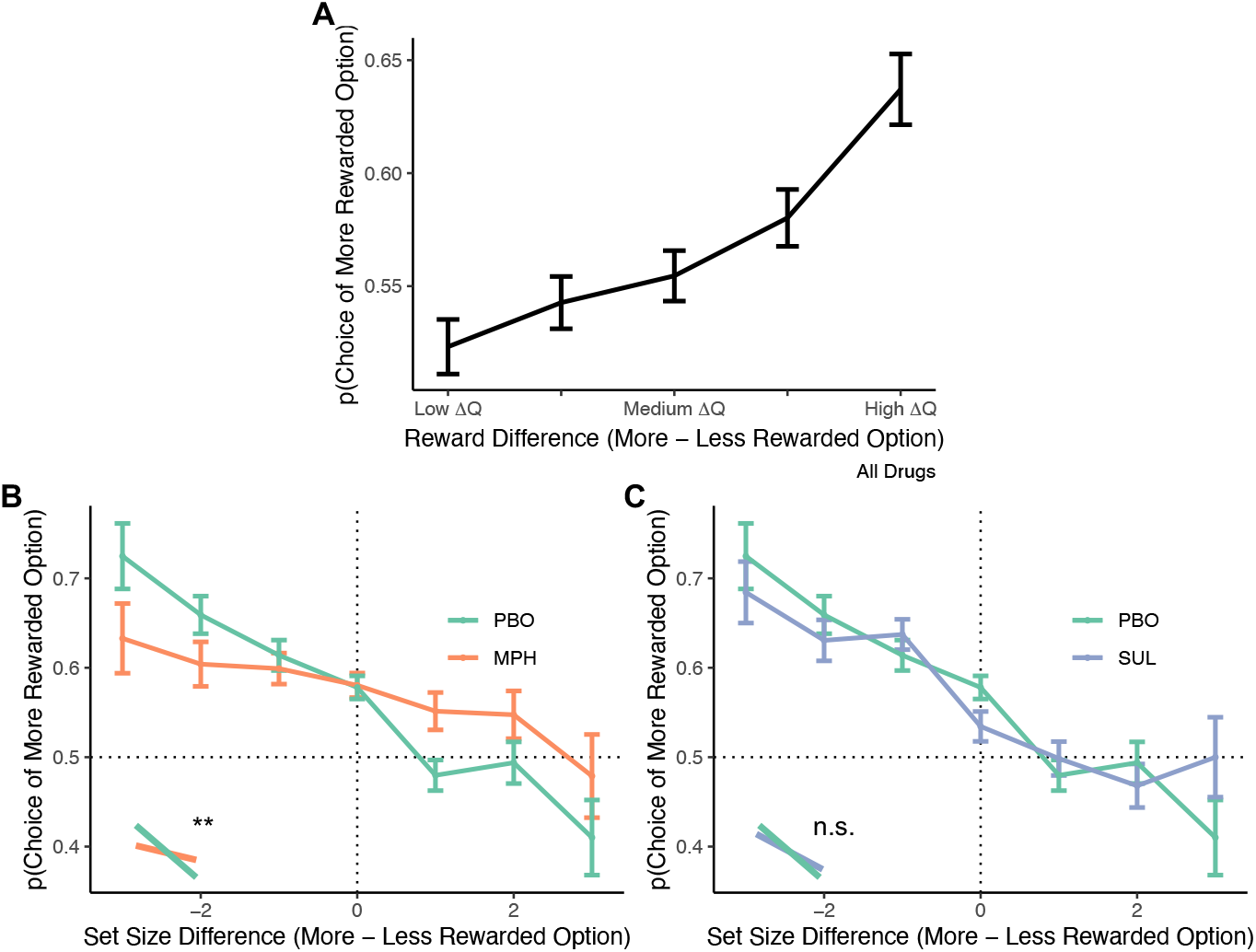
Test phase performance (selection of the option rewarded at a higher rate) as a function of the differences in **A)** actual reward statistics and **B)** and **C)** the set size of the block from which the stimulus was drawn. **A)** Participants successfully identify the more highly rewarded outcome higher than chance, and increasingly so as the difference in actual reward rates increases. **B)** and **C)** Participants are less likely to indicate an option was more highly rewarded if it came from a larger set sized block. Relative to placebo, this set size difference effect is smaller on methylphenidate and no different on sulpiride.

Also replicating prior work [39], we find that cached reward values reflect effort discounting. Specifically, controlling for the rewards people received for each stimulus during the learning phase, participants assign a lower reward value to stimuli that had been encountered in larger set sized blocks. This effect is captured by a negative effect of the difference in set sizes between the stimulus which objectively received more rewards and the stimulus which received fewer rewards (β = -.28; p = 7.1×10^−8^; Figure 6B: collapsed across differences in actual reward rates). That is, while participants track the rewards associated with each stimulus, they treat rewards received in the context of higher set sizes, and thus higher WM demands, as subjectively less rewarding. This effect has previously been interpreted as a form of effort discounting [39]. It also converges with our own prior work in which we find that participants treat higher WM demands as effort costly, requiring greater reward offers to offset these costs [18,24].

Importantly, we also find evidence that striatal dopamine blunts the discounting of rewards by the effort exerted to attain them. Specifically, the effort discounting effect of set size on rewards was significantly less on methylphenidate versus placebo (β = .12; p = .025; Figure 6B). The effort discounting effect of set size on rewards was not influenced by dopamine synthesis capacity (β = -.01; p = .85). In a different experiment conducted within the current study we find that striatal dopamine signaling increases sensitivity to reward benefits and decreases sensitivity to effort costs during decision-making about cognitive effort [24]. This result converges with the present finding that dopamine signaling can blunt the impact of effort on reward learning. Thus, by amplifying striatal dopamine signaling, methylphenidate not only increases sensitivity to benefits versus costs during action selection [24], but appears to alter how people learn about effort costs and benefits in the first place.

## Discussion

Striatal dopamine promotes corticostriatal plasticity and thereby facilitates reinforcement learning (RL) [1–4]. Numerous studies have attempted to link stronger dopamine signaling capacity to faster RL. However, this exercise is complicated by the fact that striatal dopamine may also govern the degree to which people rely on fast and flexible working memory (WM) to accomplish the same tasks [10]. Potential effects on both RL and WM implies that it is essential to control for either factor to determine the extent to which dopamine affects the other. This is true whether the goal is to evaluate the “true” rate of reinforcement learning, or the degree to which working memory contributes to learning beyond RL processes.

In this study, we employ a task which dissociates the relative contributions of RL and WM systems to learning and examine how dopamine influences each. We find that striatal dopamine signaling reflecting either dopamine synthesis capacity or dopamine drugs – can promote learning, with distinct effects on RL and WM. Specifically, higher dopamine synthesis capacity in the dorsal caudate nucleus predicts greater reliance on WM while antagonism of dopamine receptors with sulpiride reduces performance by reducing reliance on WM. By blocking dopamine reuptake and thereby amplifying dopamine signaling, methylphenidate also boosts performance in interactions with striatal dopamine synthesis capacity. Specifically, methylphenidate increases RL learning rates for people who synthesize dopamine at a higher rate and a trending interaction indicating that it tends to increase reliance on WM, particularly for those who synthesize at a lower rate.

We speculate that reliance on WM correlates with striatal dopamine synthesis capacity because the latter may shape a trait policy to rely on WM in general, in novel contexts. This could help explain why dopamine synthesis capacity may correlate with individual differences in WM capacity [26,40]. Correlations between WM capacity and dopamine synthesis capacity may obtain if study designs are sensitive to individual differences in the degree to which people rely on WM (although they will not always obtain [41]). But why does dopamine synthesis capacity promote reliance on WM? One idea is related to the emerging hypothesis that WM is effort-costly and stronger striatal dopamine signaling helps overcome those effort costs [24,27,38,42–52]. In a prior study, we showed that people are more willing to accept offers to perform more demanding WM tasks for money if they have higher dopamine synthesis capacity and on methylphenidate versus placebo [24]. Our interpretation was that striatal dopamine signaling influences both the learning and expression of cost-benefit policies governing WM allocation – making people more sensitive to performance benefits and less sensitive to effort costs.

The present findings constitute an important complement to that conclusion. Namely, while the prior study primarily examined prospective decisions about WM tasks, after effort costs had already been learned, the present study reveals that dopamine also shapes value learning about effort costs themselves. Specifically, we find that people treat rewards earned in the context of higher WM demands as subjectively less valuable – discounting rewards by increasing cognitive effort costs. Critically, we find that methylphenidate blunts this effort-discounting effect. Thus, stronger striatal dopamine signaling can make people both more willing to exert effort for tasks they have experienced in the past, and also experience the tasks as less costly when they learn about them in the first place (cf. [50]). Note that these effects of dopamine on prospective decision-making and effort cost learning complement to the proposed role of norepinephrine in energizing on-going effort expenditure [53]. Interestingly, in our present study, methylphenidate boosted reliance on WM the most for people with low dopamine synthesis capacity, mirroring our earlier finding that it also boosts explicit willingness to do cognitive effort the most among the same group [24]. The capacity of methylphenidate to blunt effort cost learning and promote willingness to exert future cognitive effort among individuals with low dopamine synthesis capacity may help account for the therapeutic effects of the drug in attention deficit hyperactivity disorder which is characterized by both pronounced working memory deficits [54] and lower dopamine synthesis capacity [55,56].

Many forms of psychopathology have been associated with aberrant rates of RL. However, our results show that it is crucial to control for the degree to which WM contributes to the learning process when trying to estimate learning rates for an RL system. Without a WM component in our learning algorithm, the learning rates for the RL system would have to be at least an order of magnitude larger to capture the rapid rates at which people acquired stimulus-response associations (in most cases one or two trials; cf. Figure 2B). Moreover, without accounting for WM contributions to the learning process, the learning rates in the RL system would have to vary by set size. Thus, both within- (e.g. set size) and between-subjects factors (e.g. WM reliance, *ρ*) would confound estimates of incremental learning rates in an RL system if WM is not accounted for. This result highlights the prospect that many prior studies showing psychopathology-linked deficits in RL may have instead found evidence of psychopathology-linked deficits in WM.

However, controlling for the contributions of WM to the learning process, we also find evidence that striatal dopamine signaling accelerates RL. Specifically, we find that methylphenidate boosts RL rates the most for people who synthesize dopamine faster in the dorsal caudate nucleus. This result mirrors prior work showing that methylphenidate promotes plasticity for reinforcement learning about rewards, albeit at the expense of specificity, especially for people with high working memory capacity [57–59] – a proxy for dopamine synthesis capacity [40]. Moreover, in another experiment conducted in the context of the current study, we find that methylphenidate increases accuracy and prefrontal BOLD signal response to reward versus punishment trials, in a reversal learning task, to a greater extent among individuals with higher dopamine synthesis capacity [60]. One explanation for these effects is that methylphenidate acts by blocking reuptake, and so it will have an even larger effect for people who synthesize and therefore release more dopamine in response to reward in the first place. The combination of greater release and reuptake blockade causes dopamine to linger in the synapse for a longer duration [61], thereby constructively amplifying LTP in response to phasic dopamine learning signals.

Although sulpiride clearly undermines performance, the mechanisms are somewhat less resolved. Sulpiride causes performance to decline both early and late in blocks suggesting it may impact both RL and WM processes. Model parameters suggest that sulpiride diminishes performance because it reduces the degree to which WM contributes to behavior. Our analyses further suggests that WM contributes less because of a faster decay of WM contents. Prior work has shown that WM and RL systems work cooperatively to accomplish learning tasks but, paradoxically, higher fidelity WM contents attenuate RL because better WM-based predictions result in smaller prediction errors [7,9]. Thus, if anything, faster-decaying, degraded WM representations should yield stronger RL and better performance late in a block. The fact that performance is also worse late in blocks suggests that sulpiride may have detrimental effects on both RL and WM systems.

While prior studies have shown a role for D2 receptors in supporting working memory function [62–64], we did not predict that antagonism of D2 receptors with sulpiride would necessarily undermine performance or reduce reliance on WM. In fact, sulpiride may strengthen rather than weaken post-synaptic dopamine signaling by binding to pre-synaptic, D2 autoreceptors, thereby releasing a break on dopamine release. In our own prior study [24], we found evidence that was consistent with the hypothesis that this dose of sulpiride strengthens post-synaptic dopamine signaling. In that study, we had independent measures (e.g. increased saccadic vigor) converging on the hypothesis that sulpiride increased postsynaptic dopamine signaling – though we are unable to make strong conclusions here. If the drug does have the same postsynaptic effects in the context of this task, we speculate that sulpiride may produce steeper effective WM decay by lowering the barrier for WM gating. If the barrier is sufficiently low, hyper-flexibility would undermine the stability of task-relevant representations across trials. Indeed, in prior studies, we have found that both the dopamine precursor levodopa [65] and the D2 agonist bromocriptine [66] can increase distractor vulnerability in cognitive control and working memory tasks. Pre-synaptic effects in our prior study helped explain why sulpiride increased willingness to perform more demanding working memory tasks by making people less sensitive effort costs in our prior study. Thus, perhaps sulpiride, when binding pre-synaptically, may both reduce sensitivity to the effort costs, and make WM less effective by amplifying decay effects.

## Conclusion

Our results support multiple, complementary mechanisms by which striatal dopamine influences task-learning. Namely, we find that individual differences in dopamine synthesis capacity correlate positively with baseline propensity to rely on working memory and working memory reliance can also be boosted for those who synthesize dopamine at a lower rate using methylphenidate. We also find that sulpiride reduces reliance on working memory, perhaps by undermining the stability of working memory contents over time. Controlling for the degree to which working memory contributes to learning processes, we also find a role of striatal dopamine on promoting plasticity for reinforcement learning. Specifically, methylphenidate boosts the rate of RL, and especially so for individuals who synthesize dopamine at a higher rate. Finally, we find evidence that pharmacological enhancement of striatal dopamine signaling can blunt effort cost learning that happens when people perform demanding tasks. This result complements prior work by showing that dopamine not only influences effort-based decision-making at the time of choice, but also by shaping the how people learn about effort in the first place.

## Methods

100 Healthy, young adult participants (ages 18—43, 50 men) were recruited from Nijmegen, The Netherlands to participate in a within-subject, double-blind, placebo-controlled study. Participants were screened to ensure that they are right-handed, Dutch-native speakers, healthy, neurologically normal, and without a history of mental illness or substance abuse. The study was part of a larger investigation of the effects of dopaminergic drugs on cognitive control. A full characterization of our participant pool, a detailed list of exclusion criteria, intake procedures, full drug administration protocol, and methods for measuring dopamine synthesis capacity as well as the full set of covariate measures and tasks for this broader study are detailed in Määttä et al., 2021 [67]. The study was approved by the regional research ethics committee (Commisssie Mensgebonden Onderzoek, region Arnhem-Nijmegen; 2016/2646; ABR: NL57538.091.16).

### General Procedure and Tasks

Participants completed five visits as part of a broader study of the effects of dopamine on cognitive control: one screening session, three pharmaco-imaging sessions with multiple tasks performed in and out of the fMRI scanner after being administered placebo, sulpiride, or methylphenidate, and a final PET session for measuring dopamine synthesis capacity. Drug session order was imperfectly counterbalanced. Consequently, 23, 15, and 10 participants took placebo on session number 1, 2, and 3, respectively, while the numbers were 12, 18, and 18 for sulpiride, and 13, 15, and 20 for methylphenidate. Given data loss and imperfect counterbalancing of drug by session order, we confirmed all inferences via hierarchical regression analyses controlling for session order.

During screening, after providing written consent, participants completed – among other tests – medical and psychiatric screening interviews, as well as tests of working memory capacity, and fluid intelligence.

Participants were asked to refrain from smoking or drinking stimulant-containing beverages the 24 hours before a pharmaco-imaging session, and refrain from using psychotropic medication and recreational drugs 72 hours before each session and cannabis throughout the experiment. At the beginning of a session, we measured baseline subjective measures, mood and affect, as well as temperature, heart rate, and blood pressure at baseline (also recorded after drug administration). Other tasks completed by participants, the results of which have been reported elsewhere, included tasks assessing sensitivity to cognitive effort costs and benefits [24,27], tasks measuring creativity [68,69], and a Pavlovian-to-instrumental transfer task [59]. Participants also completed two tasks in the fMRI scanner: one measuring striatal responsivity to reward cues and a reversal learning task [60]. Finally, after the behavioral sessions, but before the PET session, we also collected measures of depression, state affect, BIS/BAS, impulsivity, and the degree to which participants pursue cognitively demanding activities in their daily life.

Participants were administered drugs prior to the task. To accomplish double-dummy blinding participants took one drug capsule at each of two different time points: the first was either placebo or 400 mg sulpiride, while the second was either placebo or 20 mg methylphenidate. 160 minutes after taking methylphenidate or placebo (or placebo on sulpiride days), or 250 minutes after sulpiride, participants performed the RLWM task.

### Reinforcement Learning Working Memory Task

The RLWM task was presented using Psychtoolbox-3 for MATLAB. As described in the main text, participants completed two task phases: a training phase and a test phase.

In the training phase, participants were presented with stimuli in blocks of varying set sizes (between 2 and 5 stimuli in each block). Stimuli were presented one-at-a-time and participants responded with one of three button presses which were assigned at random for each stimulus. Participants were tasked with learning which of three buttons corresponded to each stimulus through trial-and-error. Stimuli were presented in pseudo-random order, for nine iterations, before switching to a new block. Participants completed between 2 and 3 blocks of each set size, also presented in pseudo-random order, and new stimuli were used for each block.

If participants responded correctly on a given trial, they were given reward feedback (+1 or +2 points probabilistically: 20/80, respectively), and if they were incorrect, they were given zero points. The number of points awarded on reward trials was not contingent on performance and, correspondingly, we found no differences between computational models fit when assuming 0,1, or 2 points for reward trials versus those fit assuming a simpler binary {1,0} for correct / incorrect trials. As such, we fit the learning algorithm (see section on “Computational modeling of behavior”, below) assuming the {1,0} binary.

At the end of the training period, participants completed a surprise test phase in which pairs of stimuli were drawn from across all blocks and participants were tasked with selecting which of each pair was rewarded at a higher rate. Participants were not given feedback about the accuracy of their response in the test phase. Because participants were assigned the task of selecting which stimuli received more points during the test phase, our analyses of the test phase assumed that participants responded based on encoding reward outcomes as either 0,1, or 2 points.

One participants’ data was excluded from analysis because their average late-block accuracy in the training phase was below 53% in all three sessions. A single session from another participant was excluded because they did not complete the training phase. In addition, three participants did not participate in the methylphenidate session, and one participant did not complete their placebo session. In total, 95 out of 100 methylphenidate, 97 out of 100 placebo, and 99 out of 100 sulpiride sessions were included in the final analysis of the training phase. In the test phase, an error with response logging meant that some sessions were excluded based on the criteria that we failed to capture participants’ responses on at least 80% of trials. Additionally, two participants were excluded based on their choice patterns which indicated that they merely alternated left / right presses on more than 75% of trials. In total, 69 out of 100 methylphenidate, 78 out of 100 placebo, and 80 out of 100 sulpiride sessions were included in the final analysis of the test phase.

For analysis of training phase accuracy, we included trials starting after the first correct iteration of each stimulus to avoid analyzing performance variance due to luck at the beginning of each block. Finally, for the test phase, we excluded trials with a response time of faster than 250 ms to avoid analyzing trials on which participants were merely guessing. The average of participants’ median response times was 1070 ms with a standard deviation of 443 ms.

Trial-wise response accuracy in both phases was modeled with fully random, Bayesian mixed effects logistic regression models using Stan for full Bayesian inference, unless otherwise indicated. The *brms* package version 2.8.0 was used to fit mixed effects regression models along with R version 3.4.3. *p*-values were calculated from the effect estimate and standard errors estimated from the upper and lower 95% confidence intervals.

For our reinforcement learning model, we adapted a variant [12] of the original model [10] which has been successfully applied to capture behavior and neural dynamics during the RLWM task in both healthy and disordered populations [8,39]. A wide range of models were fit via hierarchical maximum likelihood using the mfit toolbox (https://github.com/sjgershm/mfit) in Matlab and uniform priors for the working memory capacity, learning rate, and working memory reliance parameters. Because parameters were going to bounds for the working memory decay, punishment neglect, and testing noise parameters, we used very slightly informative priors (beta distributions with parameters *α* = 1.05 and *β* = 1.05). The model with the best BIC score was selected. Fit quality for the winning model was confirmed with posterior predictive checks comparing real data to data simulated from the model (Figure S1A-B). Parameter interpretability was confirmed via parameter recovery exercises where combinations of parameters values were selected at random from across the range of fitted values, data were simulated from each model, and models re-fit. Finally, re-fitted parameter values were compared to the original parameter values (Figure S2).

Following a previous approach [12], WM contributions were not dynamically adjusted, in the algorithm, based on the inferred reliability of WM versus RL. While this elegant approach captures dynamic adjustments to WM contributions across a block, we found that doing so reduced model recoverability. Instead, we allowed WM reliance to scale linearly with the number of unique, intervening stimuli since the last correct response for each stimulus, reasoning that WM was likely to contribute less to the extent that people encounter new, competing information since their last experience with a given stimulus. Allowing WM to contribute dynamically as a function of unique, intervening items helped account for delay effects within blocks and influenced overall reliance to WM for larger versus smaller blocks.

### PET Scanning

To measure dopamine synthesis capacity, participants completed a PET scanning session using a Siemens mCT PET-CT scanner with 40 slices, 4×4 mm in-plane voxels, and 5 mm thick slices. Prior to scanning participants received 185MBq (5mCi) F-DOPA injections into an antecubital vein. To increase [^18^F]-FDOPA concentrations, participants took 150 mg carbidopa to decrease peripheral decarboxylase activity, and 400 mg entacapone to decrease peripheral COMT activity one hour prior to injection. The 89-minute PET scan comprised four 1-minute frames, then three 2-minute frames, three 3-minute frames, and finally 14 5-minute frames.

PET images were reconstructed with weighted attenuation correction, time-of-flight correction, correction for scatter, and then smoothed with a 3 mm full-width-half-max kernel. To correct for head movement, frames were realigned to the middle frame, using SPM12. Next, images were co-registered with a structural T1-weighted MRI scan (collected in the first screening session). Presynaptic dopamine synthesis capacity was calculated as the F-DOPA influx rate (Ki; min^-1^) per voxel using the Gjedde-Patlak linear graphical analysis method for frames between 24 and 89 minutes and were referenced to signal in the cerebellum grey matter. FreeSurfer was used to segment each participants’ high-resolution anatomical MRI scans. Ki maps were normalized to MNI space and smoothed with an 8 mm full-width half-max Gaussian kernel. Finally, mean Ki values were extracted from sub-regions of the striatum, including the dorsal caudate nucleus, defined in a prior study on the basis of cortical functional connectivity patterns [70].

### Computational modeling of behavior

We hypothesized that higher dopamine signaling may influence reliance on WM and RL processes. To test these hypotheses more precisely, we adapted a learning algorithm [10,12] involving both WM and RL modules to support stimulus-response learning and fit it to behavior.

In the model, RL proceeds by tracking action values via Rescorla-Wagner style updating. Specifically, the value of a particular action *a* for a given stimulus *s* (or “policy”: Q(*s, a*)) in the RL system is updated, during learning, according to correct versus incorrect outcomes (*r*), and a learning rate parameter (*α*_*RL*_) that, following [8] is discounted (by *γ*) on incorrect trials to capture the tendency to neglect feedback about incorrect responses.

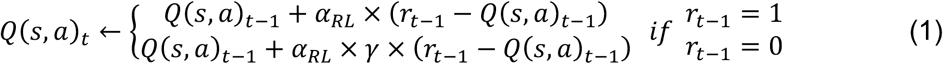

The WM system also tracks action policies (WM(*s, a*)), with an instantaneous effective learning rate.

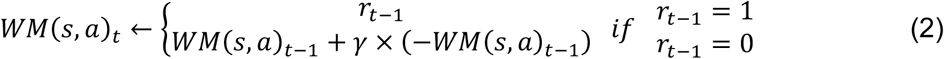

We assume that participants start each block with the belief that all actions are equally likely to be correct for a given stimulus, and so assign an initial value of *Q*(*s, a*) = 1⁄*n*_*a*_, where *n*_*a*_ is the number of possible actions (3) in this task. The WM system is subject to decay (rate: *φ*) for all stimulus-action pairs that are not observed on a given trial which then decay back towards the initial value prior.

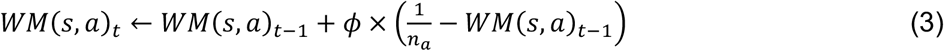

Action values from both the RL and WM systems are converted into action probabilities via the softmax function, and the respective probabilities *p*_*RL*_ and *p*_*WM*_ are combined linearly. Note that because the inverse temperature parameter can trade off with key parameters including the learning rate and WM reliance, we chose to fix the inverse temperature at a high value (*β* = 50) across all participants.

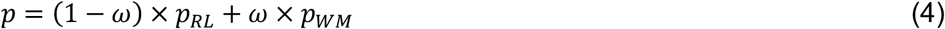

The degree to which participants rely on WM versus RL to select actions (*ω*) is defined by the parameter *ρ*, which can be thought of as a participant’s baseline propensity to rely on WM versus RL, across contexts. Overall reliance also depends on the ratio of WM capacity (*WM*_*cap*_) to the number of unique, intervening stimuli encountered since the last correct response for any given stimulus, *k*, (*n*_*delay*_,_*k*_). This modulation of *ρ* allows reliance on WM to decrease dynamically, per item, as a function of delay.

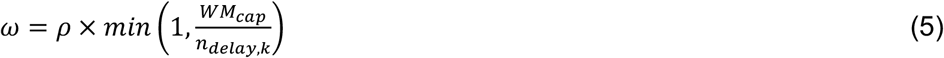

Finally, to further constrain our parameter estimates, we also leverage information about choices made during the test phase. Specifically, we included a critic which learns at the same rate as the RL actor (*α*_*RL*_), accumulating information about the value of a stimulus (averaged across all actions), *V*(*s*)_*RL*_.

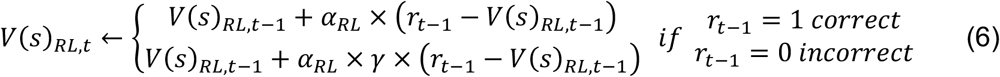

We assume that the probability that a person selects a stimulus as being rewarded the most in each pair *p* also depends on a softmax transformation (again, with fixed *β* = 50) of stimulus values to *p*_*V, RL*_ and a noise term *ν* to accommodate guessing during the test phase.

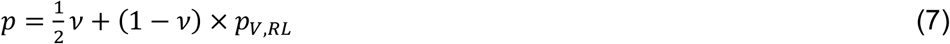

We fit our model by maximum likelihood. The fitted model captures qualitative effects of set size and iteration number (Supplementary Figure S1A). It also captures delay, early versus late trials and the interactions of these variables with dopamine variables including individual differences in dopamine synthesis capacity, and drug effects (Supplementary Figure S1B,C). The model not only provides an excellent fit to the data but is also recoverable, indicating that individual participant-level parameter estimates are reliable and interpretable (Supplementary Figure S2). Prior to evaluating how parameters are altered by dopamine synthesis capacity and drugs, we removed participants with outlier values (more than ± 2 standard deviations from the mean) for parameters of interest to reduce the likelihood that any single extreme and unlikely estimate exerts too much leverage on the evaluation process. This resulted in the removal of 5 participants from the methylphenidate session (5.3% of participants), 4 from the placebo session (4.2%) and 2 from the sulpiride session (2.0%).

## Data availability

The datasets generated during and/or analyzed during the current study are available in the Radboud Data Repository, https://doi.org/10.34973/wn51-ej53.

## Author Contributions

**Andrew Westbrook:** Conceptualization, Methodology, Formal Analysis, Writing – Original Draft. **Ruben van den Bosch**: Investigation, Formal Analysis, Visualization, Writing – Review & Editing. **Lieke Hofmans**: Investigation, Writing – Review & Editing. **Danae Papadopetraki:** Investigation, Software, Writing – Review & Editing. **Jessica I. Määttä**: Methodology, Investigation, Data Curation, Writing – Review & Editing. **Anne G E Collins**: Software, Conceptualization, Writing – Review & Editing. **Michael Frank:** Conceptualization, Supervision, Formal Analysis, Writing-Reviewing and Editing. **Roshan Cools:** Conceptualization, Supervision, Formal Analysis, Writing-Reviewing and Editing.

