## Supplemental Info for "Striatal Dopamine Can Enhance Learning, Both Fast and Slow, and Also Make it Cheaper"

### Supplement

*Bayesian mixed effects model of training phase accuracy*

The following mixed effects logistic regression was fitted to trial wise accuracy across all participants and drug conditions. All effects had both random and fixed effects terms. The model was fit using the *brms* package version 2.8.0 in R. Note that the logistic regression model is fit based on delay modeled as $-1/n_d$ where $n_d$ is the number of trials since the last previous correct iteration, and all numeric predictors are z-scored.

**Supplemental Table S1**

| | Trial-wise Training Phase Accuracy | | |
|---|---|---|---|
| *Predictors* | *Log-Odds* | *CI* | *p* |
| Intercept | 3.56 | 3.42 – 3.72 | **<0.001** |
| Set Size ($n_s$) | -0.15 | -0.24 – -0.06 | **0.001** |
| Delay ($n_d$) | -0.05 | -0.14 – 0.04 | 0.75 |
| DA Synth. Capacity (DA) | 0.17 | 0.02 – 0.32 | **0.026** |
| Methylphenidate (MPH) | 0.41 | 0.24 – 0.57 | **<0.001** |
| Sulpiride (SUL) | -0.27 | -0.41 – -0.13 | **<0.001** |
| Previous Correct Iterations (pCor) | 0.58 | 0.49 – 0.67 | **<0.001** |
| Session Number (Sess) | 0.10 | 0.04 – 0.15 | **<0.001** |
| $n_s$ * $n_d$ | -0.28 | -0.35 – -0.21 | **<0.001** |
| $n_s$ * DA | -0.10 | -0.019 – -0.01 | **0.029** |
| $n_d$ * DA | 0.04 | -0.04 – 0.13 | 0.35 |
| $n_s$ * MPH | 0.02 | -0.11 – 0.14 | 0.75 |
| $n_s$ * SUL | 0.03 | -0.08 – 0.14 | 0.65 |
| $n_d$ * MPH | -0.07 | -0.20 – 0.06 | 0.30 |
| $n_d$ * SUL | -0.11 | -0.23 – -0.00 | **0.047** |
| DA * MPH | -0.07 | -0.23 – 0.09 | 0.40 |
| DA * SUL | -0.01 | -0.15 – 0.13 | 0.90 |
| $n_s$ * pCor | 0.16 | 0.08 – 0.24 | **<0.001** |
| DA * pCor | -0.01 | -0.10 – 0.08 | 0.84 |
| MPH * pCor | 0.20 | 0.06 – 0.33 | **0.004** |
| SUL * pCor | -0.04 | -0.15 – 0.07 | 0.49 |

| | | | |
|---|---|---|---|
| $n_d$ * pCor | 0.15 | 0.08 – 0.23 | **<0.001** |
| $n_s$ * $n_d$ | -0.28 | -0.35 – -0.21 | **<0.001** |
| $n_s$ * $n_d$ * DA | -0.08 | -0.14 – -0.01 | **0.016** |
| $n_s$ * $n_d$ * MPH | -0.02 | -0.12 – 0.09 | 0.72 |
| $n_s$ * $n_d$ * SUL | -0.01 | -0.11 – 0.08 | 0.85 |
| $n_s$ * DA * MPH | 0.03 | -0.10 – 0.15 | 0.65 |
| $n_s$ * DA * SUL | 0.06 | -0.04 – 0.17 | 0.27 |
| $n_d$ * DA * MPH | -0.08 | -0.21 – 0.04 | 0.21 |
| $n_d$ * DA * SUL | -0.02 | -0.13 – 0.09 | 0.73 |
| $n_s$ * DA * pCor | 0.03 | -0.05 – 0.11 | 0.47 |
| $n_s$ * MPH * pCor | -0.04 | -0.17 – 0.09 | 0.56 |
| $n_s$ * SUL * pCor | -0.06 | -0.17 – 0.05 | 0.29 |
| DA * MPH * pCor | 0.02 | -0.11 – 0.15 | 0.78 |
| DA * SUL * pCor | 0.01 | -0.09 – 0.11 | 0.86 |
| $n_d$ * DA * pCor | -0.06 | -0.13 – 0.02 | 0.90 |
| $n_d$ * MPH * pCor | 0.07 | -0.05 – 0.20 | 0.76 |
| $n_d$ * SUL * pCor | -0.00 | -0.11 – 0.10 | 0.99 |
| DA * MPH * pCor | 0.02 | -0.11 – 0.15 | 0.78 |
| DA * SUL * pCor | 0.01 | -0.09 – 0.11 | 0.86 |
| $n_d$ * DA * pCor | -0.06 | -0.13 – 0.02 | 0.90 |
| $n_d$ * MPH * pCor | 0.07 | -0.05 – 0.20 | 0.76 |
| $n_d$ * SUL * pCor | -0.00 | -0.11 – 0.10 | 0.99 |
| $n_s$ * $n_d$ * DA * MPH | 0.05 | -0.05 – 0.15 | 0.33 |
| $n_s$ * $n_d$ * DA * SUL | 0.10 | 0.01 – 0.19 | **0.029** |
| $n_s$ * DA * MPH * pCor | -0.02 | -0.15 – 0.10 | 0.77 |
| $n_s$ * DA * SUL * pCor | -0.05 | -0.15 – 0.06 | 0.36 |
| $n_d$ * DA * MPH * pCor | 0.06 | -0.06 – 0.18 | 0.33 |
| $n_d$ * DA * SUL * pCor | 0.11 | 0.00 – 0.21 | **0.036** |
| $N_{id}$ | 92 | | |
| Observations | 77699 | | |

*Bayesian mixed effects model of test phase accuracy*

The following mixed effects logistic regression was fitted to trial wise selection of the stimulus which was rewarded at the higher rate in the test phase across all participants and drug conditions. All effects had both random and fixed effects terms. The model was fit using the *brms* package version 2.8.0 in R. Note that the logistic regression model is fit to trials with reaction times slower than 0.25 seconds, and all numeric predictors are z-scored.

**Supplemental Table S2**

| | Trial-wise Testing Phase Accuracy | | |
|---|---|---|---|
| *Predictors* | *Log-Odds* | *CI* | *p* |
| Intercept | 0.08 | 0.02 – 0.14 | **0.009** |
| Value Difference ($\Delta V$) | 0.31 | 0.23 – 0.38 | **<0.001** |
| Mean Set Size ($\bar{n}_s$) | 0.03 | -0.01 – 0.08 | 0.19 |
| Mean Value ($\bar{V}$) | -0.00 | -0.05 – 0.05 | 0.97 |
| DA Synth. Capacity (DA) | 0.01 | -0.05 – 0.08 | 0.78 |
| Methylphenidate (MPH) | -0.01 | -0.09 – 0.06 | 0.81 |
| Sulpiride (SUL) | -0.05 | -0.13 – 0.02 | 0.19 |
| Set Size Difference ($\Delta n_s$) | -0.28 | -0.38 – -0.18 | **<0.001** |
| Trial Number | 0.03 | -0.00 – 0.06 | 0.057 |
| Session Number | 0.01 | -0.02 – 0.05 | 0.59 |
| $\Delta V$ * $\bar{n}_s$ | 0.06 | 0.00 – 0.11 | **0.028** |
| $\Delta V$ * $\bar{V}$ | -0.01 | -0.09 – 0.08 | 0.83 |
| $\Delta V$ * DA | 0.02 | -0.06 – 0.10 | 0.14 |
| $\bar{n}_s$ * DA | -0.01 | -0.06 – 0.05 | 0.73 |
| $\bar{V}$ * DA | -0.04 | -0.10 – 0.02 | 0.19 |
| $\Delta V$ * MPH | 0.09 | -0.05 – 0.23 | 0.21 |
| $\Delta V$ * SUL | 0.02 | -0.09 – 0.14 | 0.75 |
| $\bar{n}_s$ * MPH | -0.01 | -0.08 – 0.06 | 0.79 |
| $\bar{n}_s$ * SUL | -0.01 | -0.08 – 0.05 | 0.78 |
| DA * MPH | 0.03 | -0.05 – 0.11 | 0.47 |
| DA * SUL | -0.02 | -0.10 – 0.07 | 0.66 |
| $\bar{V}$ * $\Delta n_s$ | 0.01 | -0.06 – 0.08 | 0.79 |

| | | | |
|---|---|---|---|
| DA * $\Delta n_s$ | -0.01 | -0.13 – 0.06 | 0.85 |
| MPH * $\Delta n_s$ | 0.12 | 0.01 – 0.22 | **0.025** |
| SUL * $\Delta n_s$ | 0.03 | -0.06 – 0.12 | 0.52 |
| $\Delta V$ * $\overline{n_s}$ * $\overline{V}$ | 0.04 | -0.02 – 0.10 | 0.19 |
| $\Delta V$ * $\overline{n_s}$ * DA | 0.02 | -0.04 – 0.09 | 0.56 |
| $\Delta V$ * $\overline{V}$ * DA | 0.04 | -0.03 – 0.12 | 0.30 |
| $\overline{n_s}$ * $\overline{V}$ * DA | 0.01 | -0.04 – 0.07 | 0.73 |
| $\Delta V$ * $\overline{n_s}$ * MPH | -0.02 | -0.13 – 0.09 | 0.73 |
| $\Delta V$ * $\overline{n_s}$ * SUL | -0.09 | -0.18 – 0.00 | 0.47 |
| $\Delta V$ * $\overline{V}$ * MPH | 0.11 | -0.01 – 0.24 | 0.084 |
| $\Delta V$ * $\overline{V}$ * SUL | 0.01 | -0.10 – 0.14 | 0.88 |
| $\overline{n_s}$ * $\overline{V}$ * MPH | 0.08 | -0.00 – 0.16 | 0.053 |
| $\overline{n_s}$ * $\overline{V}$ * SUL | 0.07 | 0.00 – 0.15 | 0.061 |
| $\Delta V$ * DA * MPH | 0.03 | -0.13 – 0.17 | 0.71 |
| $\Delta V$ * DA * SUL | -0.05 | -0.17 – 0.06 | 0.40 |
| $\Delta V$ * $\overline{n_s}$ * MPH | 0.03 | -0.04 – 0.10 | 0.41 |
| $\Delta V$ * $\overline{n_s}$ * SUL | -0.05 | -0.13 – 0.01 | 0.16 |
| $\overline{V}$ * DA * MPH | 0.04 | -0.03 – 0.13 | 0.33 |
| $\overline{V}$ * DA * SUL | -0.00 | -0.09 – 0.08 | 0.93 |
| $\overline{V}$ * DA * $\Delta n_s$ | 0.04 | -0.02 – 0.11 | 0.23 |
| $\overline{V}$ * MPH * $\Delta n_s$ | -0.02 | -0.13 – 0.09 | 0.73 |
| $\overline{V}$ * SUL * $\Delta n_s$ | -0.02 | -0.12 – 0.08 | 0.71 |
| DA * MPH * $\Delta n_s$ | -0.09 | -0.20 – 0.03 | 0.13 |
| DA * SUL * $\Delta n_s$ | -0.05 | -0.16 – 0.07 | 0.40 |
| $\Delta V$ * $\overline{n_s}$ * $\overline{V}$ * DA | -0.05 | -0.12 – 0.02 | 0.16 |
| $\Delta V$ * $\overline{n_s}$ * $\overline{V}$ * MPH | -0.01 | -0.10 – 0.08 | 0.84 |
| $\Delta V$ * $\overline{n_s}$ * $\overline{V}$ * SUL | -0.07 | -0.15 – 0.02 | 0.11 |
| $\Delta V$ * $\overline{n_s}$ * DA * MPH | -0.02 | -0.10 – 0.06 | 0.64 |
| $\Delta V$ * $\overline{n_s}$ * DA * SUL | -0.01 | -0.10 – 0.08 | 0.84 |
| $\Delta V$ * $\overline{V}$ * DA * MPH | 0.01 | -0.12 – 0.12 | 0.88 |

| | | | |
|---|---|---|---|
| $\Delta V * \bar{V} * DA * SUL$ | -0.05 | -0.19 – 0.08 | 0.48 |
| $\bar{n}_s * \bar{V} * DA * MPH$ | 0.01 | -0.09 – 0.09 | 0.84 |
| $\bar{n}_s * \bar{V} * DA * SUL$ | -0.05 | -0.14 – 0.03 | 0.25 |
| $\bar{V} * DA * MPH * \Delta n_s$ | 0.00 | -0.12 – 0.12 | 0.99 |
| $\bar{V} * DA * SUL * \Delta n_s$ | -0.09 | -0.20 – 0.01 | 0.093 |
| $\Delta V * \bar{n}_s * \bar{V} * DA * MPH$ | 0.07 | -0.04 – 0.17 | 0.19 |
| $\Delta V * \bar{n}_s * \bar{V} * DA * SUL$ | 0.05 | -0.06 – 0.15 | 0.36 |
| $N_{id}$ | 81 | | |
| Observations | 24522 | | |

*Learning algorithm – posterior predictive checks and parameter recoverability*

A key test of model quality is whether an algorithm can recapitulate behavioral patterns. To test this, we simulated data from our fitted model (Figure S1). The resulting simulations reveal that the model captures key features of the data including sensitivity to iteration number, set size, dopamine synthesis capacity, early versus late trials, and delay.

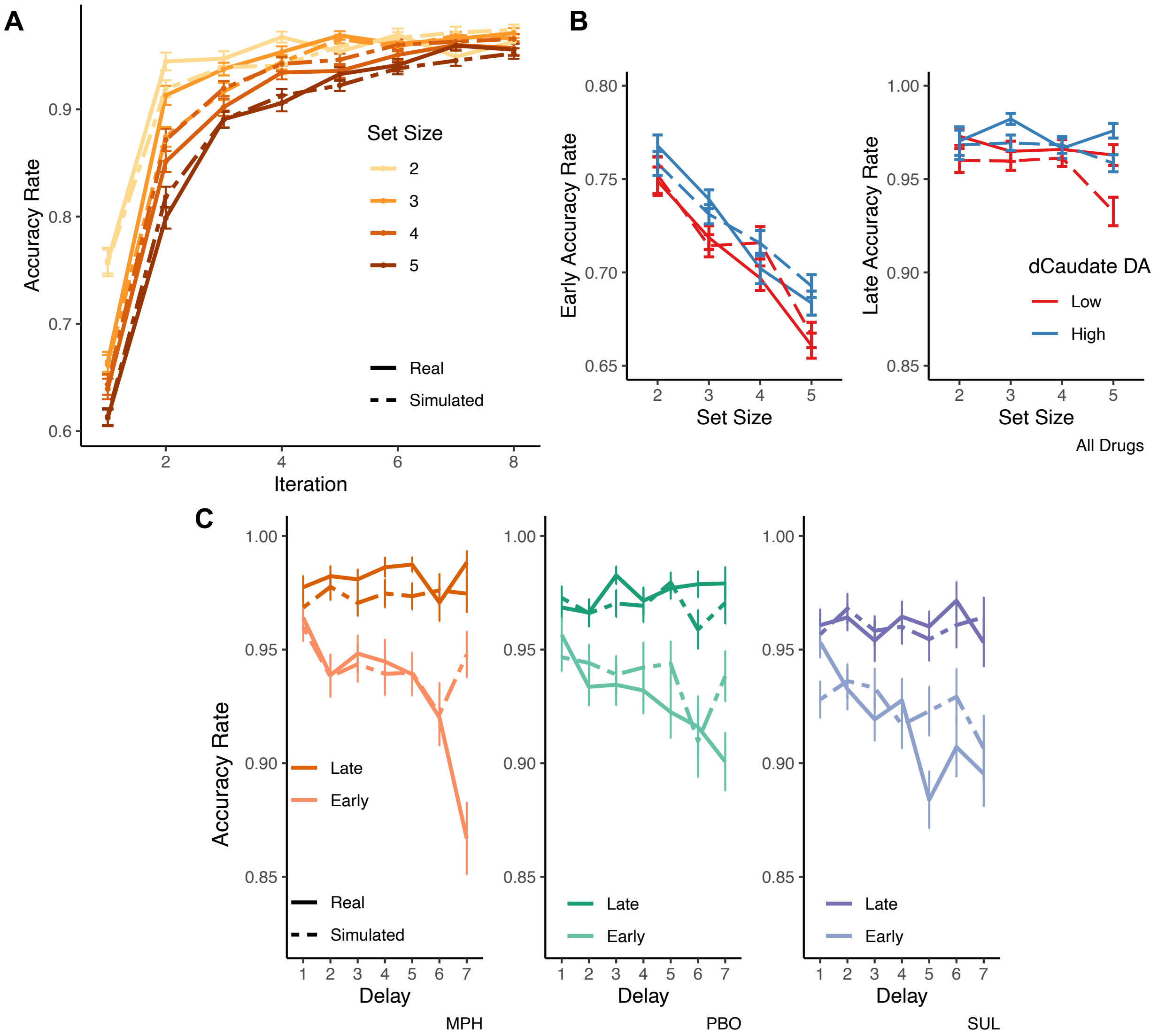


**Figure S1.** Model fit to behavior. The fitted model captures key effects including **A)** iteration number and set size and **B)** dopamine synthesis capacity and set size effects which are present early but not late in a block, and **C)** drug effects across sessions as well as the effects of delay, which are present early, but not late in a block.

While our estimated parameters – the RL learning rate ($\alpha_{RL}$) and the initial WM weighting ($\rho$), in particular – converge with model-independent analyses of behavior, the ability to draw inferences from fitted parameters depends on whether parameter estimates are recoverable. That is, inferential capacity depends on whether one can trust that fitted parameters are an accurate reflection of their true values, given the model. One way to test this is to simulate artificial data from randomly chosen parameter value combinations (where parameters values are randomly chosen from across the range of values estimated from real data), and then fit the model to the behavior generated by these simulated agents. If the fitted values of the model match the randomly chosen value of each agent, this implies that, given the model, we can trust that parameters reliable and interpretable – i.e., that they are *recoverable*.

As revealed by the strong correlations between fitted and simulated parameter values, key parameters, including both $\alpha_{RL}$, and $\rho$, are recoverable, with high fidelity (Figure S2). Some parameters, e.g. the testing phase noise, appear to not be recoverable, however this did not undermine the recoverability of key terms.

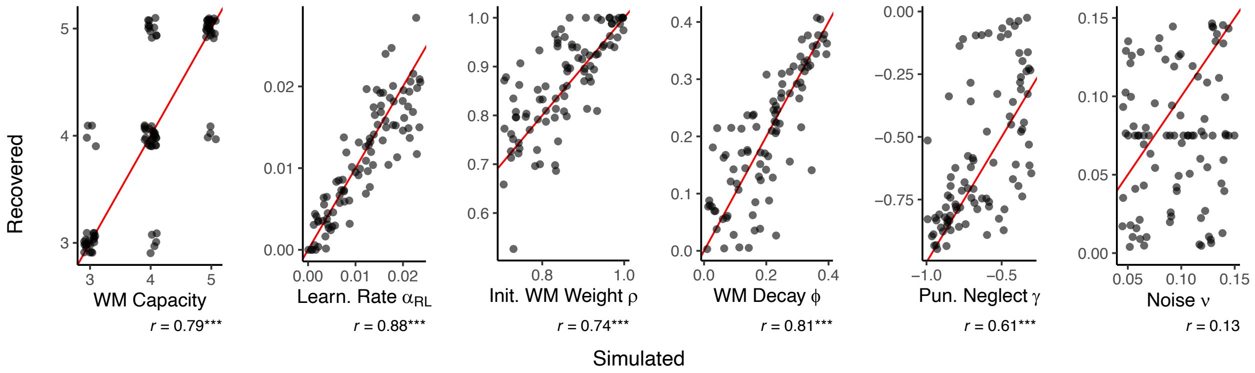


**Figure S2.** A comparison of simulated and recovered parameter values for simulated agents reveals that key parameters are recoverable in our model.
